## Supplementary figures and images for "Framework for analyzing MAE-derived immunopeptidomes from cell lines with shared HLA haplotypes"

### Supplementary Figure 1

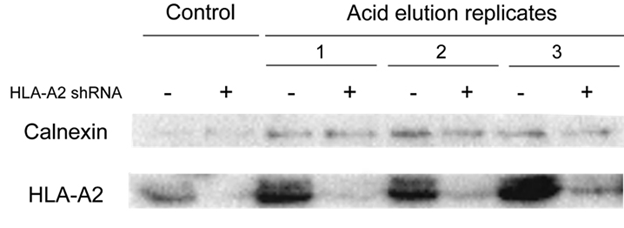

### Supplementary Figure 2

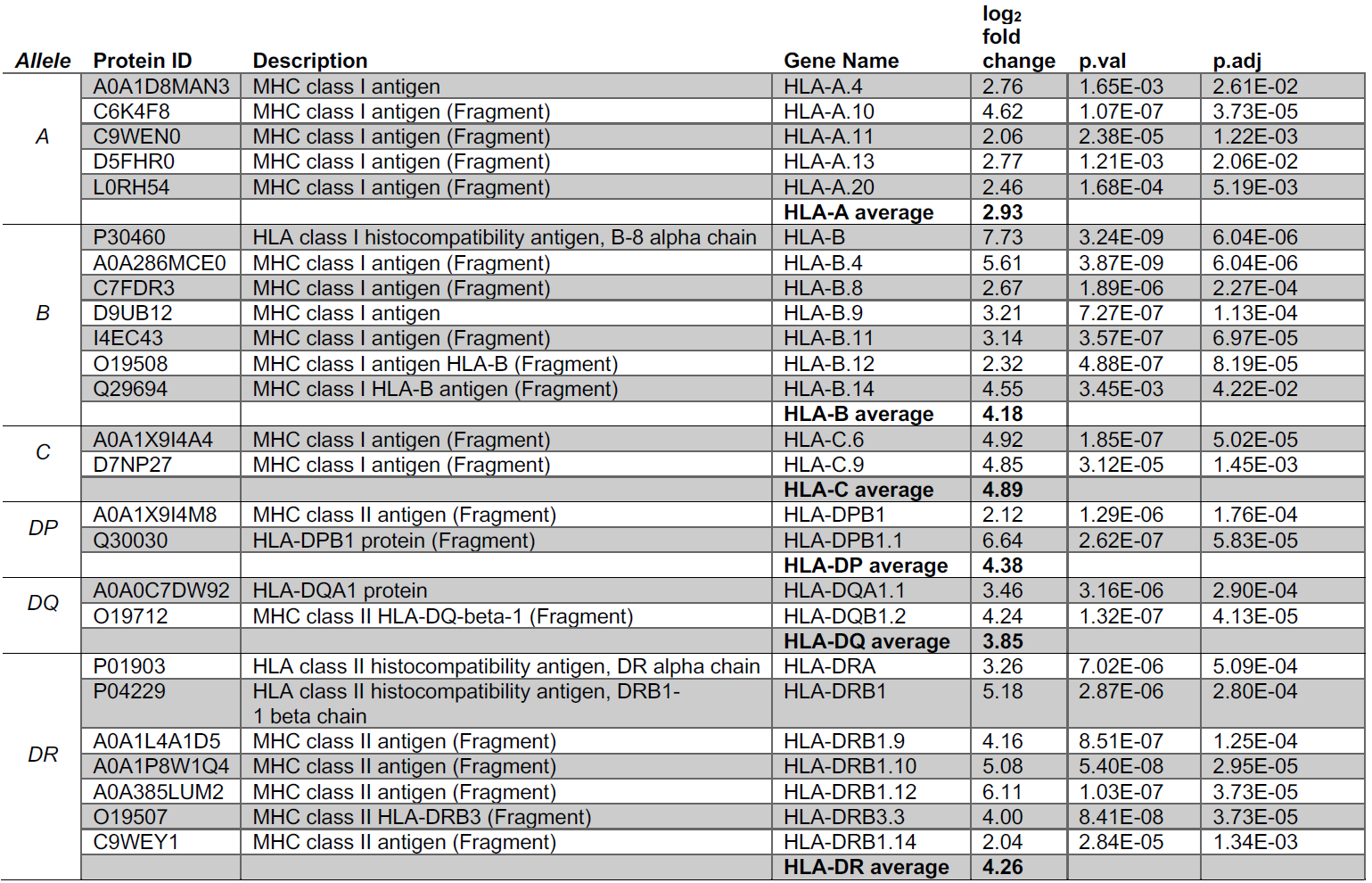

### Supplementary Figure 3

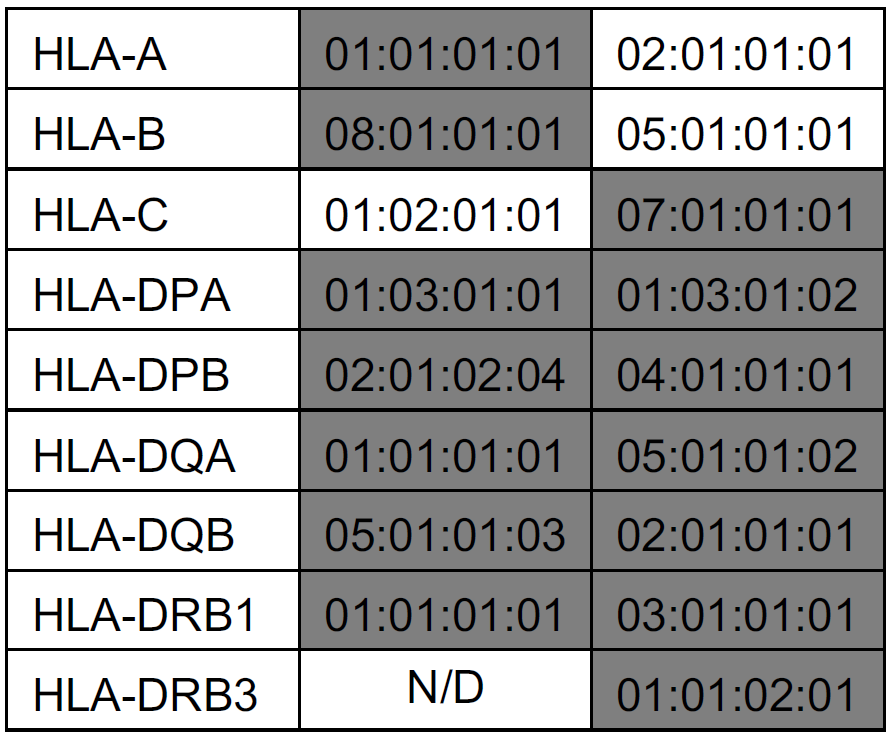

### Supplementary Figure 4

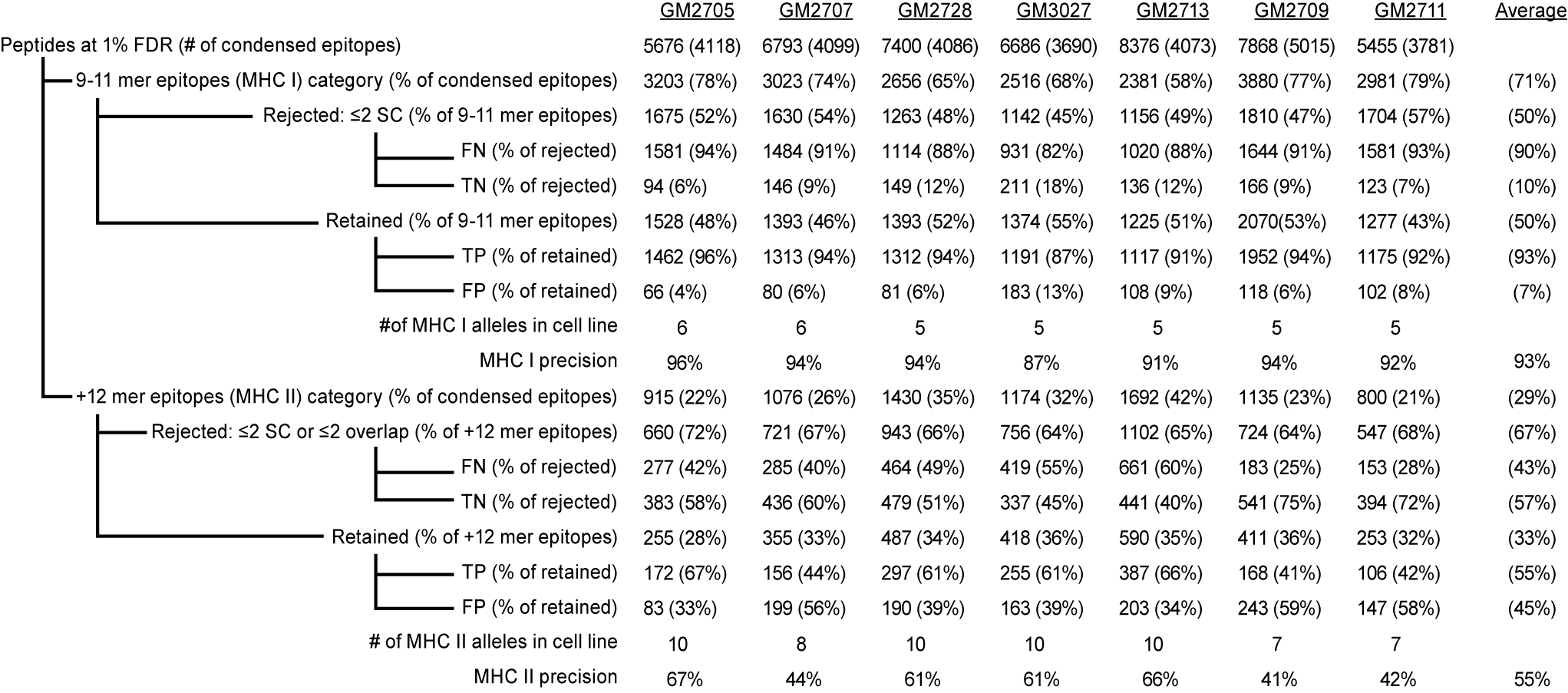

### Supplementary Figure 5

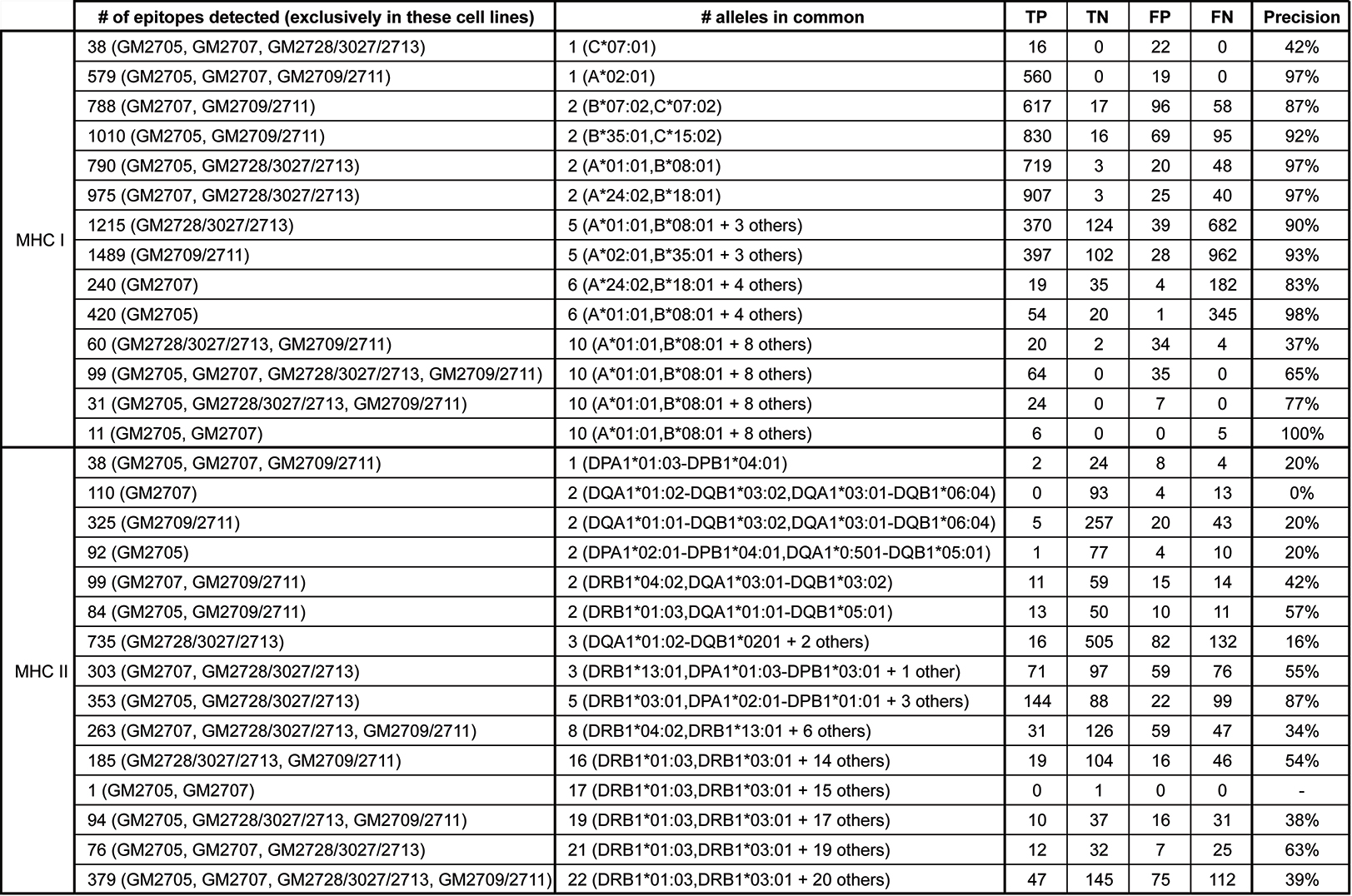

### Supplementary Figure 6

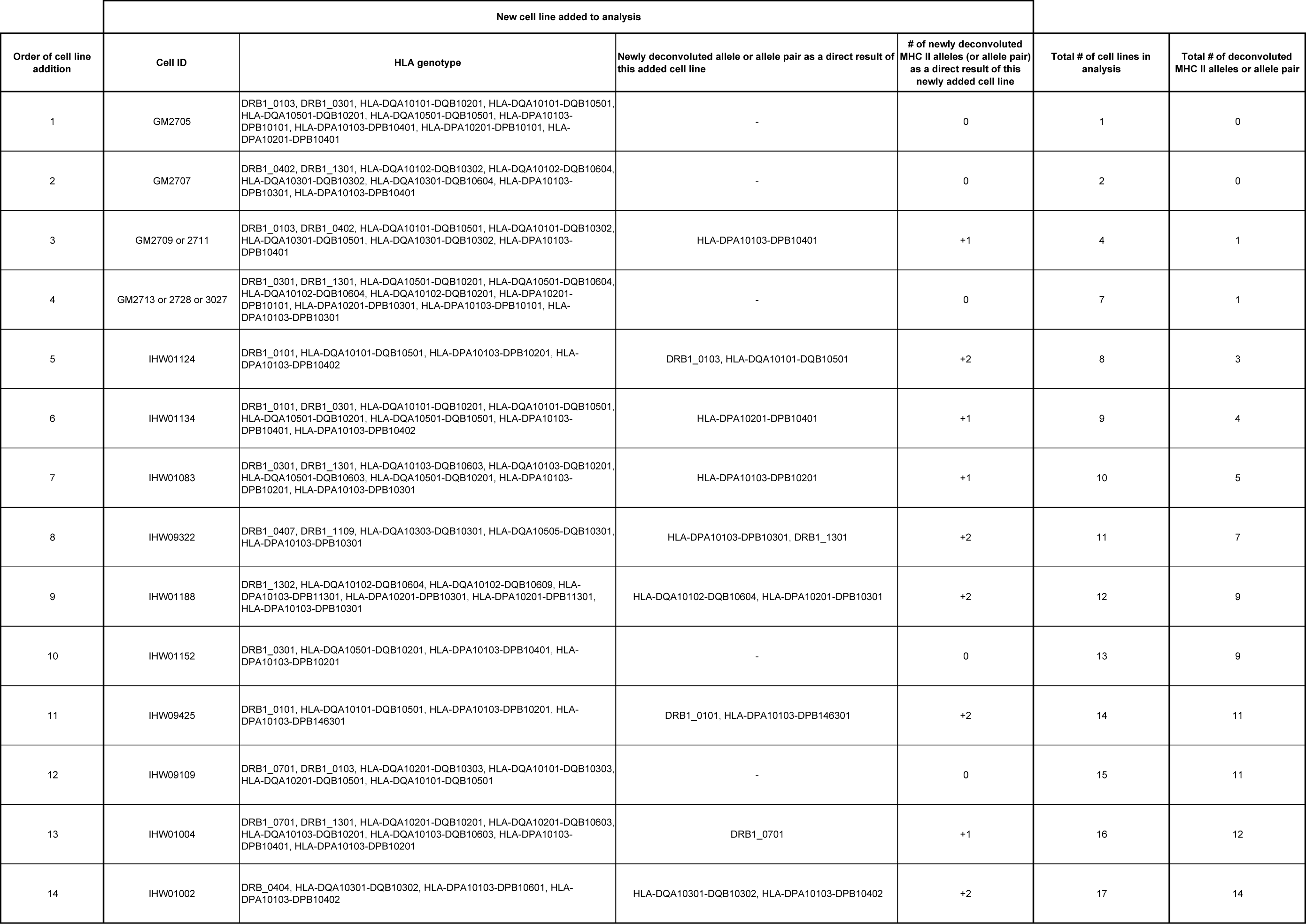

### Supplementary File 1

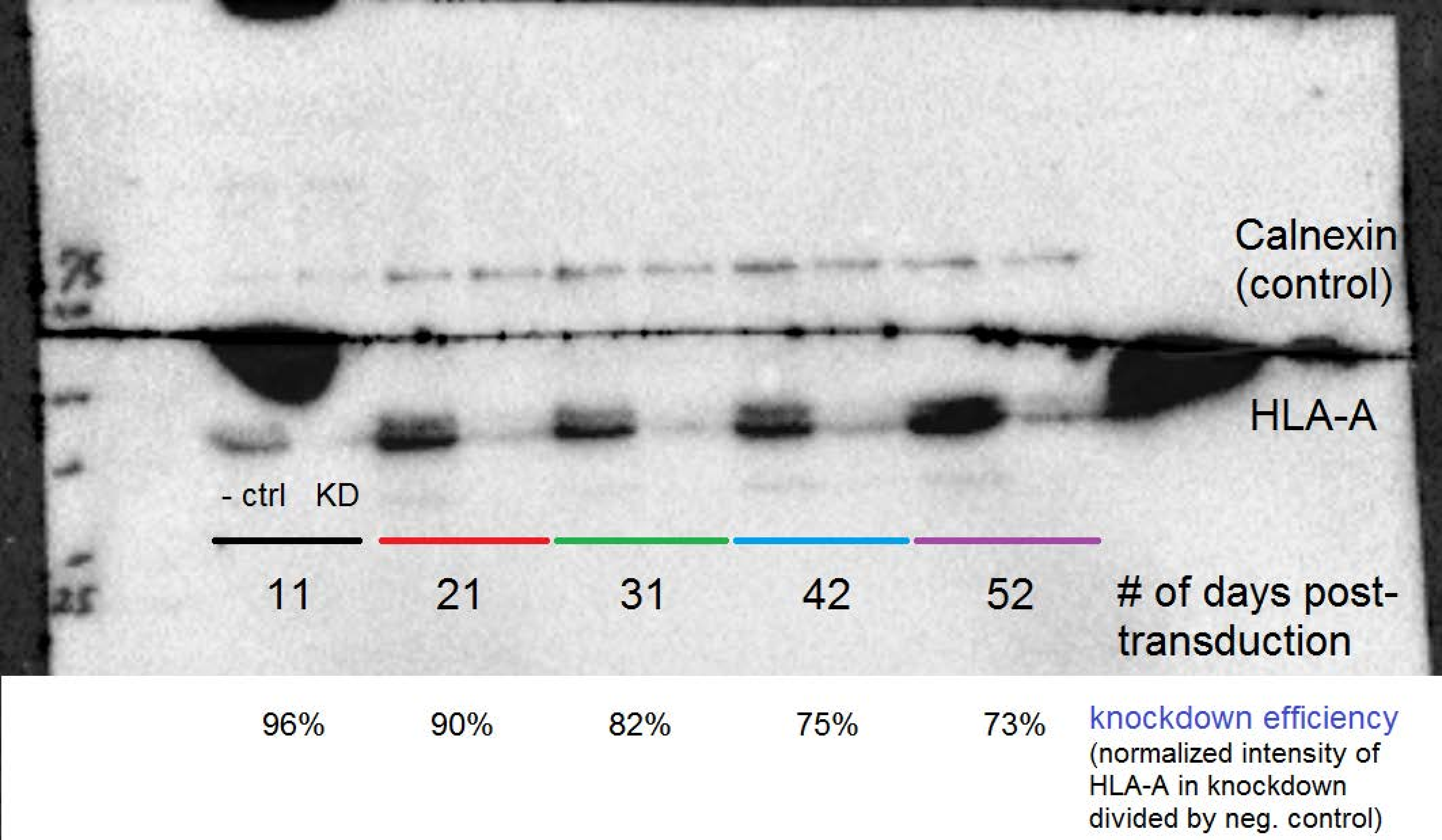
